# Spontaneous dimensional reduction and novel ground state degeneracy in a simple chain model

**DOI:** 10.1101/2021.03.13.435258

**Authors:** Tatjana Škrbić, Trinh Xuan Hoang, Achille Giacometti, Amos Maritan, Jayanth R. Banavar

## Abstract

Chain molecules play a key role in the polymer field and in living cells. Our focus is on a new homopolymer model of a linear chain molecule subject to an attractive self-interaction promoting compactness. We analyze the model using simple analytic arguments complemented by extensive computer simulations. We find several striking results: there is a first order transition from a high temperature random coil phase to a highly unusual low temperature phase; the modular ground states exhibit significant degeneracy; the ground state structures exhibit spontaneous dimensional reduction and have a two-layer structure; and the ground states are assembled from secondary motifs of helices and strands connected by tight loops. We discuss the similarities and notable differences between the ground state structures (we call these PoSSuM - Planar Structures with Secondary Motifs) in the novel phase and protein native state structures.

---

Materials made up of polymer chains are ubiquitous in everyday life and in industry. Here we study a simple model of a chain with tuned interactions, which yields very unusual behavior of the ground state conformation(s) of the chain. Intriguingly, even though the chain lives in three dimensional space, it sacrifices exploring all three dimensions and spontaneously becomes a two layer structure in order to benefit from the maximal number of contacts. The system is therefore Euclidean two-dimensional in its ground state and should not be confused with the fractal dimension of two adopted by a random walk. Furthermore, this layered structure exhibits strands and helices that are able to interchange with each other resulting in a huge ground state degeneracy. Quite remarkably, this behavior is robust on changing all but one parameters of the model (this crucial parameter needs to be held fixed to maintain the interchangeability of helices and strands). Finally, our results are *not* artifacts of finite size effects.

We consider a chain [1, 2] of *N* spheres with diameter *σ* tethered to each other in a railway train topology with a fixed bond length *b*. In what follows, all length scales are measured in units of the bond length *b*, which we set equal to 1 without loss of generality. A chain is inherently anisotropic because of tethering while an individual sphere is isotropic. In order to avert this spurious symmetry, we consider two modifications. First, adjacent spheres overlap [3, 4, 5] - *σ* is allowed to be larger than 1. Second, side spheres [6, 7, 8, 9, 10, 11, 12] of diameter *σ*_*s*_ are attached (tangent to the main chain sphere in the negative normal direction in the local Frenet frame) to each of the main chain spheres except the first and the last. Other than the adjoining main chain spheres along the chain, none of the main chain and side spheres is allowed to overlap. We impose a pairwise attractive interaction between the main chain spheres within the range of attraction *R*, and separated by at least 4 along the sequence, with an energy scale of ϵ (set equal to 1). We ascribe a distinct weaker energy of *x =* –1/2 for a pair of main chain spheres separated by exactly 3 spheres along the sequence and within the range of interaction *R*. This tuning yields the very special highly degenerate ground states assembled from building blocks of helices and strands. There is no attractive interaction for main chain spheres with sequence separation less than 3. All energies (including the temperature) are measured in units of the depth of the attractive square well potential ϵ.

The coordinate system we employ is shown in Figure 1. For fixed bond length, a chain conformation is specified by two angles *θ* and *μ. θ* is a measure of bond bending, a straight conformation has *θ = π*. Two distinct kinds of bending energy penalties have been commonly employed in the literature: an energy cost proportional to cos^2^ (*θ*/2); or zero cost when *θ* > *θ*_0_ and infinite cost otherwise [13, 14]. Here we use a third approach and make the simplification of fixing *θ* = *θ*_0_ resulting in *θ* no longer being a variable of the model but just a parameter. This can be thought of as *θ* being pinned at its minimum allowed value due to the tendency for compactness and yields a freely rotating chain [1, 2]. The angle between successive binormals, *μ*, is the second angular coordinate and is the dihedral or torsional angle. For simplicity, we do not incorporate a torsional rigidity energy, conventionally chosen to be proportional to sin^2^(*μ*/2) [15], or any other explicit energy dependence on *μ*. The parameters of the model are *σ, σ*_*s*_, *R, x*, and *θ*_0_. The PoSSuM phase is centered around *σ* = 4/3, *σ*_*s*_ = ⅔, R = 8/5, *x =* – ½, and *θ*_0_ *=* 97°. The phase is robust to small variations (of the order of 10%) in these parameter values with one exception. As we shall see, the rich non-trivial degeneracy of ground state conformations requires *x* to be – 1/2.

**Figure 1.**
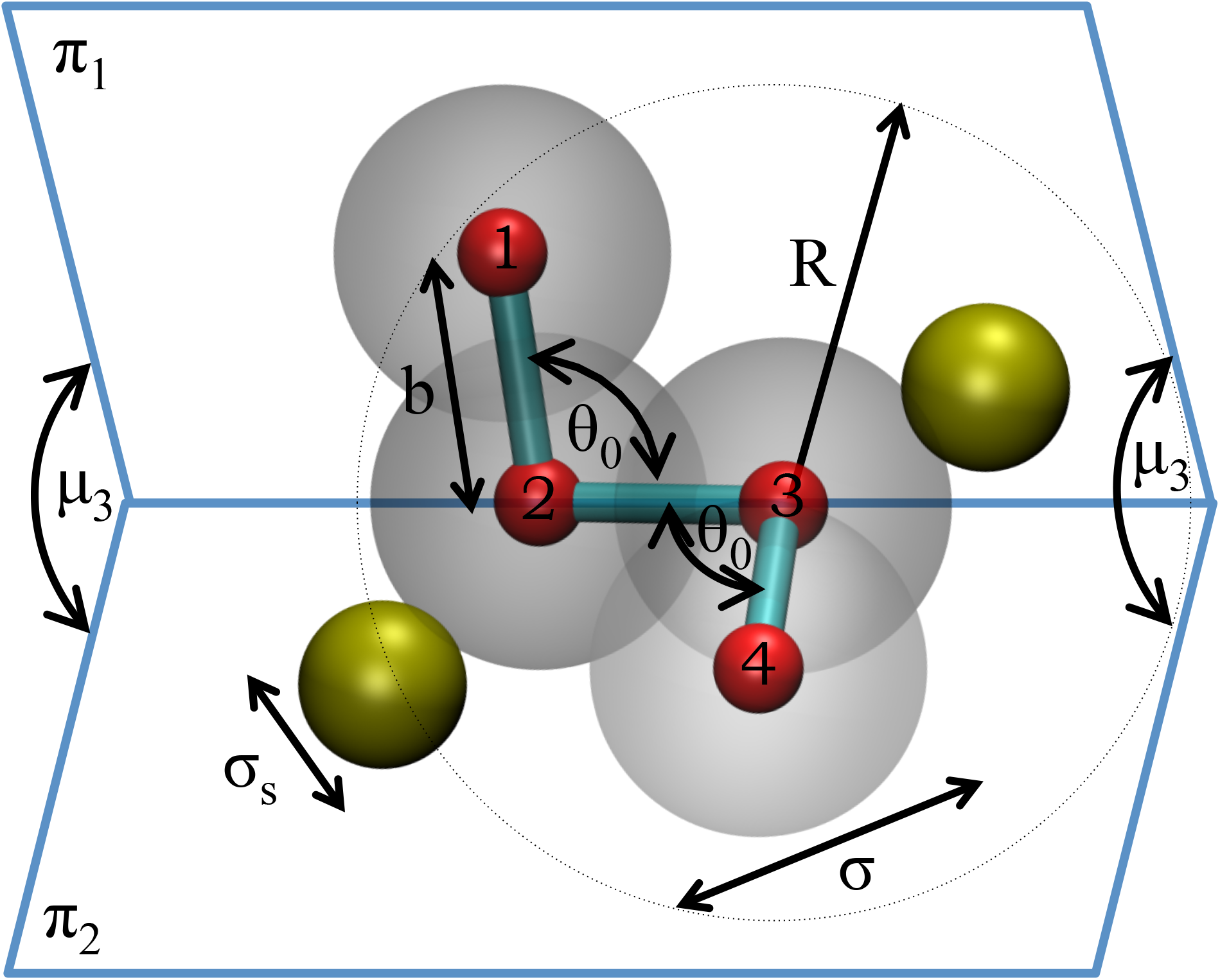
Coordinate system used in our study. Here we show 4 consecutive main chain spheres along the chain. The bond length *b* = 1 sets the length scale of the model. The main chain spheres (shown in grey with centers marked by little red spheres) have a diameter *σ* larger than *b* leading to an overlap of successive spheres. The yellow side spheres of diameter *σ*_*s*_ are attached to the main chain spheres tangentially in the negative normal direction. The angles between successive bonds are held constant at a value *θ*_0_. The range of attractive interactions is denoted by *R. μ*_*3*_ is the dihedral angle between the planes (π_1_ and π_2_) defined by sphere centers (1,2,3) and (2,3,4), respectively. It is also the angle between successive binormal vectors in a Frenet coordinate system at sphere centers 2 and 3.

The high temperature phase is a random coil and exhibits the familiar Flory scaling [1, 2] of the end-to-end distance and radius of gyration as a function of chain length (≈ *N*^0.6^). We have verified this with simulations for *N* up to 1024 (results not shown). At low temperatures, the chain adopts a compact conformation to maximize the number of attractive main sphere contacts, while respecting the chain connectivity and steric interactions. Akin to a periodic crystal, the simplest structured conformations of a chain arise from a repeat of the *μ* values along the chain. For the state point above, any uniform *μ* ≥ *μ*_*min*_ *=* 36.45° yields clash-free chains of indefinite length. A repeat structure with *μ* = *μ*_*min*_ results in the most tightly wound helix with the maximum number of local contacts - a sphere in the interior of the helix *i* has 4 contacts with spheres separated in sequence by –4 –3 +3, and +4, yielding a total attractive energy of –3 units per interior sphere (Figure 2, Panel I). The same 4 contacts remain until the *μ* value of 45.285°. One obtains a two-dimensional strand, when *μ* is close to *π* (Figure 2, Panel I). The strand does not have any within-strand contacts and has zero attractive energy per interior sphere. We note the key observation, made in the protein context [16, 17], that sterics inhibits structural hybrids and promotes structures with repeat *μ* values. Indeed, we find in simulations that a random equal choice of helix-strand *μ*-values in a chain of length 40 yields fewer than 7% of viable chains with no steric clashes.

**Figure 2.**
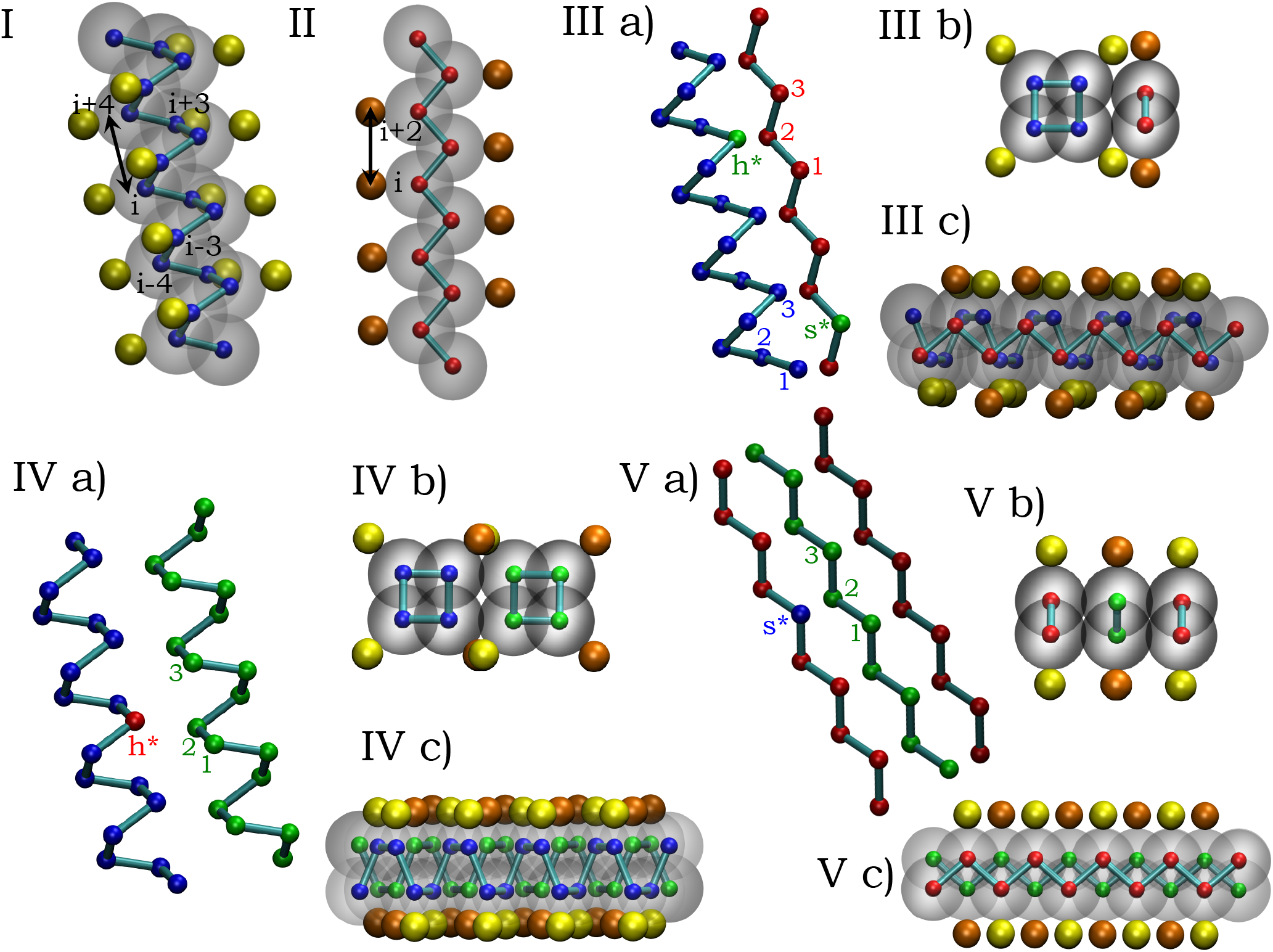
Sketches of ideal assembly of secondary motifs. Panels I and II show snapshots of an individual helix and strand with perfect commensurability attributes. The model has *θ*= *θ\** = 98.213° and the optimal values of *µ* = *µ*\* = 41.410° and 180° for the helix and strand respectively. There is commensurability both in the number of main chain spheres per repeat unit (4 for the helix and 2 for the strand) as well as equality of the lengths of the repeat units (helix (*i,i* + 4) distance = strand (*i,i* + 2) distance = 1.512 in units of the bond length). The small spheres outline the backbone of the chain, the large grey spheres are the main chain spheres (diameter = 4/3), the little yellow (for helix) (orange, for strand) spheres are the side spheres (diameter = 2/3) pointed in the negative normal direction. A solitary strand has no local contacts, whereas the local energy score per interior helix main chain sphere is –3 arising from energies of –1, – 1/2, –1/2, and – 1 for 4 contacts of sphere *i* with spheres *i* – 4,*i* – 3,*i* + 3 *i* + 4, respectively. Panels III, IV, and V show the perfect fit of a helix with a strand; two helices of opposite chirality; and three strands. The three sub-panels show three views (front, top, and side) of each of these assemblies, in particular, revealing the two layer structure. Panels III (a), IV (a), and V (a) show a trace of the backbone of the secondary motifs (without main chain or side spheres) and the three non-local contacts made by each internal sphere of the secondary motif. For a strand, the total number of non-local contacts is 6 (3 from one side and 3 from the other), whereas for a sphere in a helix, there are four local contacts (yielding a favorable energy of –3 - recall *x* = –1/2) and three non-local contacts resulting in a net favorable energy of –6. Indeed, each of the interior spheres of any of these assemblies has a net energy of –6 yielding the highly degenerate ground state. Our analysis of the PoSSuM phase in this paper is carried out with *θ*= 97° in the vicinity of *θ*\*. The flexibility in the *θ* and *µ* angles allows an excellent match of the ground state PoSSuM structures with the perfect commensurability structures shown here.

Armed with the insight that individual helices and strands are building blocks of the PoSSuM ground state structures, we proceed to work out their harmonious packing. We seek commensurability of the repeat structures - the pitch of the helix and the distance between the (*i,i* + 2) spheres or corresponding pitch in a strand, which is equal to *2b* sin(*θ* /2); and the number of spheres per turn in a helix and the number of spheres per turn in a strand, equal to 2. One can readily work out the conditions for perfect commensurability: 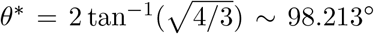 for our model with *μ*\* = 41.410° for the helix and 180° for the strand. For these choices, the special helix has exactly 4 main chain spheres per turn (with perfect commensurability with the strand) and its pitch (the *i,i* + 4 distance) exactly matches the (*i,i* + 2) distance or the corresponding pitch of the planar strand. Figure 2 (Panels III-V) illustrates the ideal packing of secondary motifs that yield a two layer idealized structure. Helices of opposing chiralities tend to pack much better than helices of the same chirality, which cannot make the same number of contacts without creating steric clashes between side spheres (not shown). It is important to note that this ideal commensurate phase point lies within the basin of the PoSSuM phase. A single helix has an energy of – 3 corresponding to 4 intra-helical contacts (recall our choice of x = – 1/2) and each sphere in the helix has exactly three inter-helical contacts (be it with a single partner helix of opposite chirality or a strand) yielding a net energy per sphere of *−*6. single strand has no attractive energy on its own but accumulates a favorable energy of *−*6 per sphere with a contribution of −3 from each of its two partners, be it another strand or a helix.

The exact degeneracy of structures in the PoSSuM phase is broken when *x* deviates from the value of *−*1*/*2. We can readily work out the ground state phases surrounding the PoSSuM phase on varying the model parameters. Too large an attraction range leads to a globular phase, whereas the converse yields a sheet phase because the inter-helical contacts are disrupted. Too large a main chain sphere size results in steric clashes whereas the converse leads to disruption of the intra-helical contacts, again leading to the sheet phase. Too small a side-sphere size yields a globule phase whereas too large a side-sphere size disrupts inter-helical contacts with the sheet again emerging as the winning phase. Finally, too small a *θ* angle does not allow for a helix with 4 spheres per turn thereby promoting the sheet phase. The sheet phase is also the phase of choice for too large a *θ* angle because of the disruption of intra-helical contacts - the distance between the main chain spheres *i* and *i* + 4 becomes prohibitively large. In this way, the PoSSuM phase is seen to be nestled between the globular phase and the sheet phase at low temperatures, thereby conferring on it the sensitivity associated with being in the vicinity of a phase transition. The robustness of the sheet phase is due to the ability to place strands at an optimal distance from each other with no major issues pertaining to side-sphere clashes. The PoSSuM phase exists in a Goldilocks window of parameter space characterized by a delicate combination of steric constraints both from the main and side spheres, the optimal range of attraction and especially the fine-tuning of the attractive interaction, as well as the fixed bond bending angle allowing for commensurability. Interestingly, the PoSSuM phase exists only when the main chain spheres overlap thereby removing the spurious spherical symmetry.

We now turn to a brief description of our computer simulations before presenting the results. The extensive search for ground state configurations was performed using Wang-Landau microcanonical Monte Carlo (MC) simulations [18] with no low energy cut-off. The density of states g(E) that are visited along the simulation was iteratively built by filling consecutive energy histograms. The acceptance probability was chosen to promote moves toward less populated energy states thus providing for increasing flatness of energy histograms with the length of the simulation. The density of states g(E), using which the thermodynamics was calculated, was obtained employing cut-off values for energy histograms within 2% of the pre-determined value of the ground state energy of the system. In all cases 28-30 levels of iterations were carried out with a flatness criterion in each iteration of at least 80%, ensuring convergence of the results [19, 20, 21]. The set of MC moves were modified to maintain *θ* constant and included both local-type moves, such as single-sphere crankshaft, reptation, and end-point moves, as well as non-local type moves, such as the pivot [22].

Figure 3 shows a gallery of low energy conformations (for *N* = 80) in the PoSSuM phase along with evidence for domain formation in simulations of *N* = 160. There are deviations from the optimal energy of *−*6 per interior sphere due to turns which connect the secondary elements together as well as significant edge effects occurring at the boundaries. A remarkable feature of the gallery is the distinct topologies of the degenerate ground state structures. The degeneracy is further enhanced by the possibility of a coordinated conversion of a helix to a strand and vice-versa while maintaining the energy. The PoSSuM phase and our model is distinct from that used in an earlier study that identified an elixir phase of matter [11, 12]. The previous model did not have a fixed *θ* angle. However moderate *θ* angles were promoted because of an *i − i* + 2 attractive interaction. Also, *x* was set equal to 1 in that model favoring the helix over the sheet.

**Figure 3.**
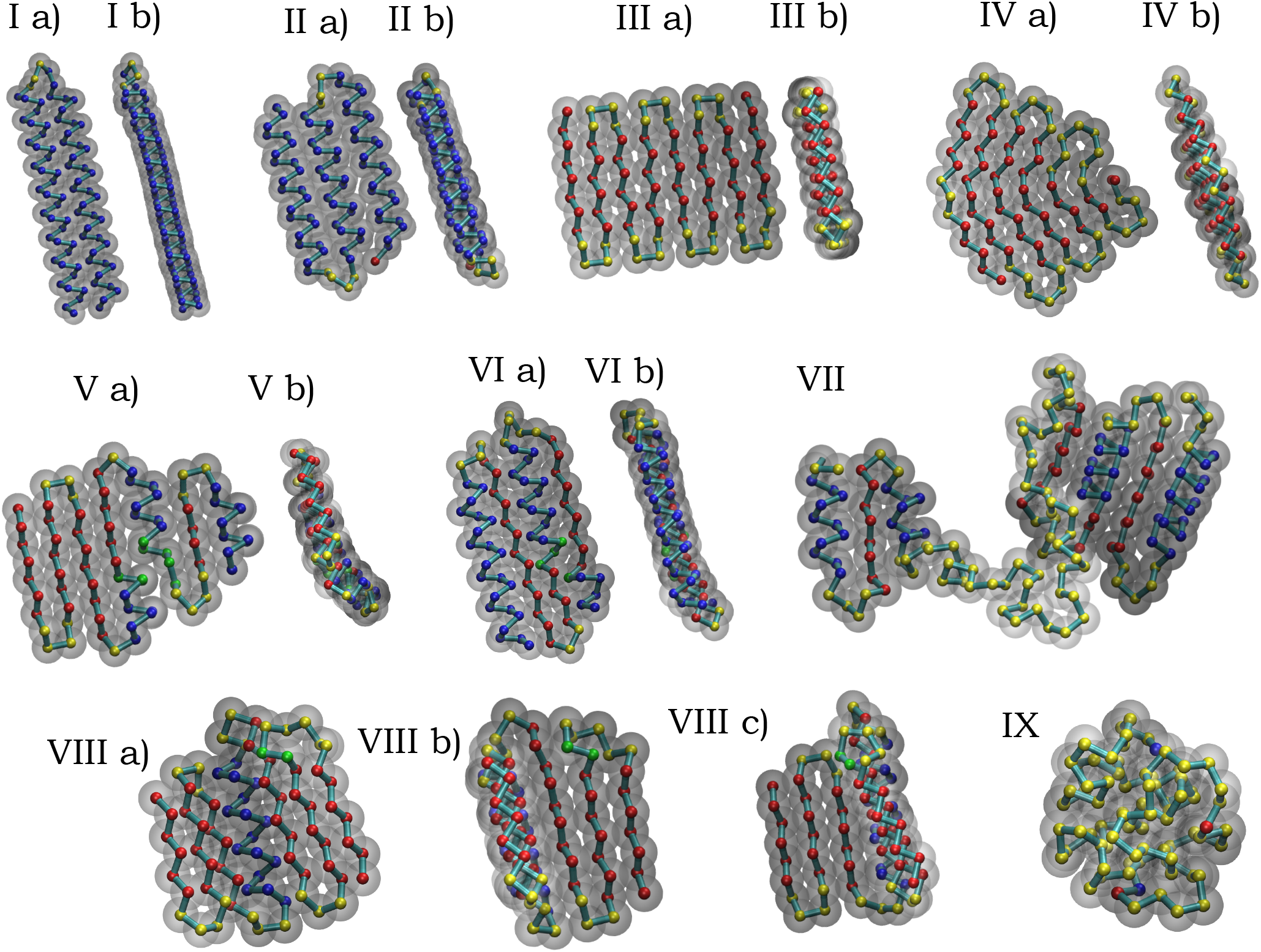
Gallery of conformations. All the panels depict conformations of chain length *N* = 80 except for VII, which shows domain formation for a *N* = 160 chain. Panel IX is a globular conformation for the case of a chain with no side spheres. The presence of side spheres results in nearly degenerate PoSSuM conformations of two helices (I), three helices (II), a sheet assembled from strands (III), a curved sheet (IV), conformations which include a switch between a helix and strand (V and VI), and a pivotal helix that links two sheets at right angles to each other (VIII). For several panels, two or three perspectives are shown to illustrate the tight packing and the layered structure. The grey circles depict the main chain spheres. The side spheres are not shown. The little circles lie at the centers of the main chain spheres. As a guide to the eye, we color a helix sphere center blue, a strand sphere center red, a turn sphere center yellow and a hybrid sphere center in the vicinity of a helix-strand switch green. These assignments are made based on *μ* values, the number of main chain spheres per turn, and the number of local and non-local contacts.

Figure 4 presents data pertaining to the nature of the phase behavior of the PoSSuM model. Panel (a) is a plot of the specific heat versus the temperature for three different chain lengths. The specific heat peak grows approximately linearly with *N* as expected for a first order phase transition [20, 23, 24]. The inset of the figure shows the canonical energy *P* (*E, T*)\, which exhibits the characteristic bimodal shape at a first order transition. Panel (b) shows a histogram of *µ* values for three different temperatures, one in the high temperature phase, another in the vicinity of the transition and the third in the low temperature phase. At low temperatures, the peaks occur at the *µ*-values of the helix, the strand, and the turns. There is a signal of the formation of secondary motifs in the histogram of *µ* angles even at a temperature more than 50% larger than the transition temperature. Panel (c) shows the layered structure of the PoSSuM ground states. The absence of sharp layering is readily accounted for because the layer separation depends on the secondary motifs adjacent to each other. The side spheres lie on the opposite ends of the layers and prevent attractive interactions with a putative third layer and growth in the third dimension. Finally, Panel (d) is the spatial configuration associated with a ground state gallery of the globular phase obtained by taking the PoSSuM model and eliminating the side-spheres.

**Figure 4.**
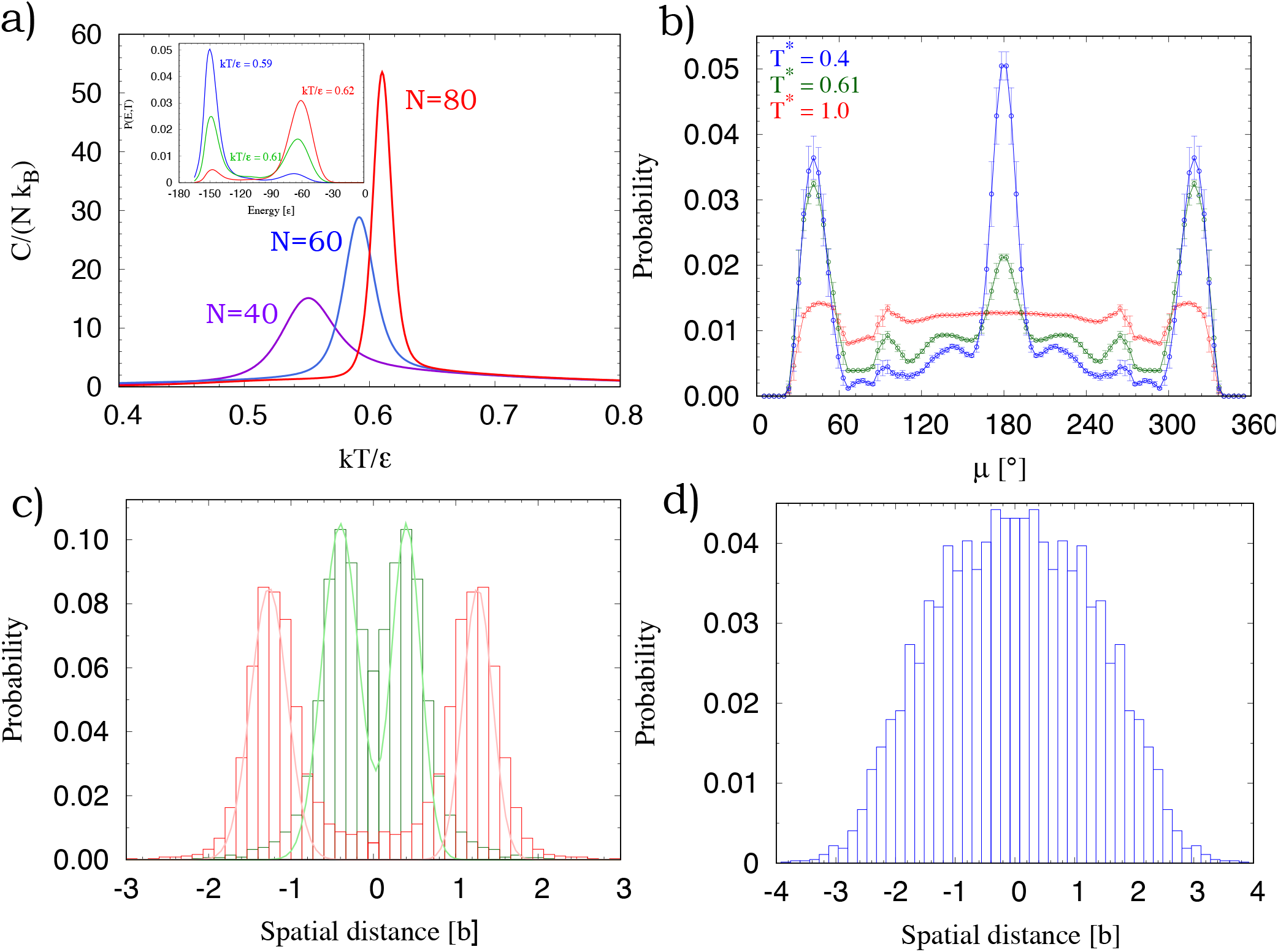
Temperature dependence of the PoSSuM model. *a*) Plot of the specific heat as a function temperature for three chain lengths. The peak sharpens and increases approximately linearly with *N* suggesting that the transition is first order. The inset is a plot of the distribution of the canonical energy for *N* = 80 for three temperatures in the vicinity of the transition. *b*) The distribution of *µ* at three temperatures. At a temperature of 1.0, there is already a signature of incipient secondary motifs. The three peaked structure at the temperature of 0.4 underscores the presence of secondary motifs. *c*) The two layered structure of the PoSSuM ground states is shown by the green histogram fitted as the sum of two Gaussians. The red histograms indicate the spatial spread of the side spheres. In order to determine the direction perpendicular to the layers, we used the eigenvector corresponding to the smallest eigenvalue of the moment of inertia matrix. The graph is an average over 40 distinct ground state PoSSuM conformations of *N* = 80. *d*) is the corresponding plot for the globular phase that is obtained when the side chains are removed from the model. Note the lack of layering and the higher spread of the main chain spheres.

Figure 5 shows the energy as a function of MC time in a single long run at a temperature equal to approximately 98% of the transition temperature. It shows multiple switching between the high temperature and low temperature phases as expected at a first order transition. Furthermore the low energy structures (we show 20 of these visited in just the single run) are very well-formed modular structures made up of helices and strands and having energies within 10% of the lowest energy conformation obtained in detailed Vang-Landau computer simulations [18]. Finally, the PoSSuM phase is not a finite size effect. In the thermodynamic limit, it is straightforward to deduce that the geometries of the degenerate ground states are still essentially planar with a thickness of two layers with both the length and width scaling as 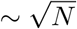.

**Figure 5.**
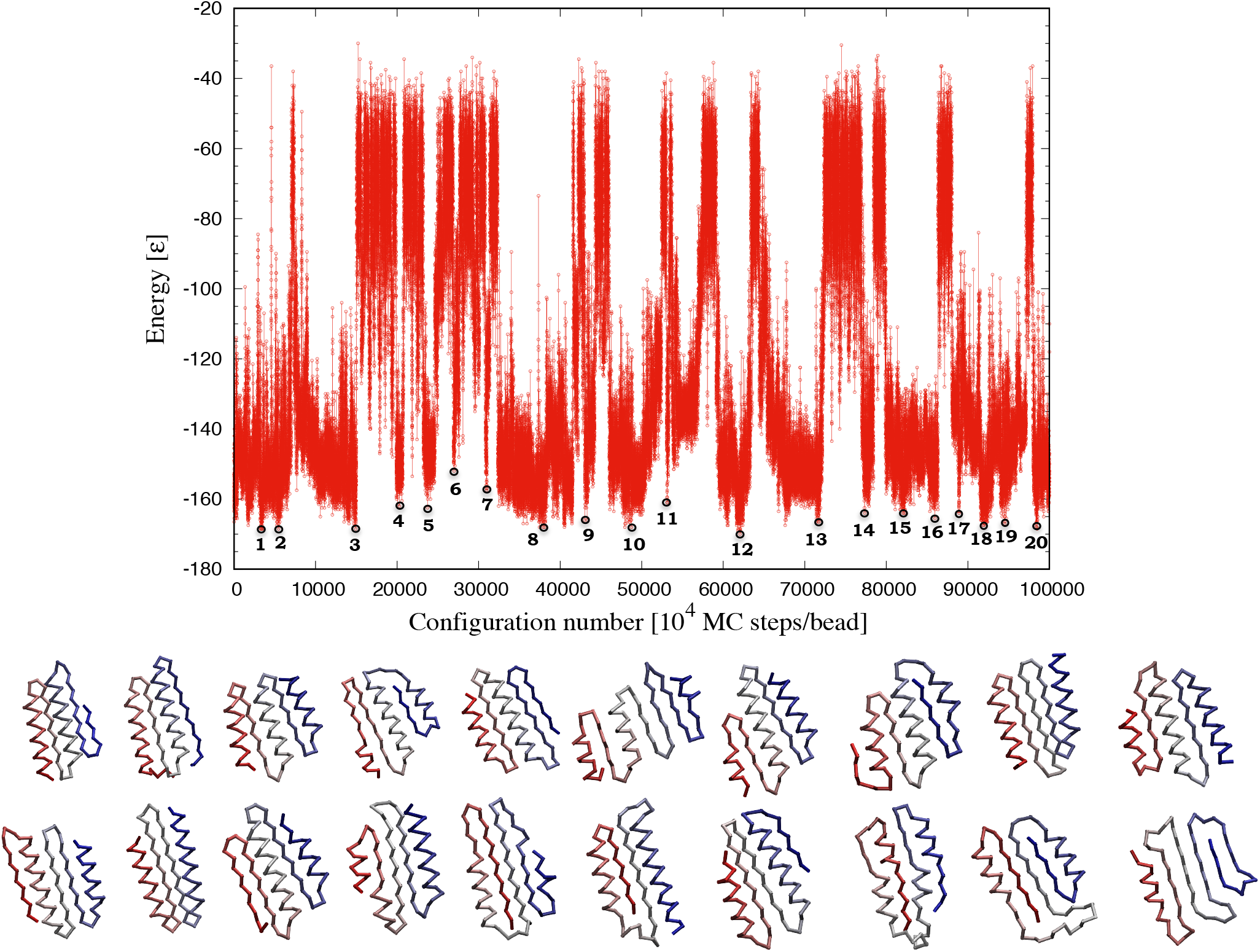
A single constant temperature Monte Carlo run for *N* = 80 at a reduced temperature of 0.595, around 2% below the transition temperature. The energy is plotted as a function of Monte-Carlo time and shows many switches between the unfolded state and the PoSSuM state. Twenty well-formed nearly degenerate PoSSuM configurations with distinct topologies are shown at the bottom of the figure in the same order as their appearance in the Monte-Carlo run.

Even though our model has features reminiscent of native structures of proteins [25, 26, 27, 28] (the modular building blocks of helices and strands arising from the presence of side-chains is one striking commonality), there are essential differences. Proteins are made up of twenty types of naturally occurring amino acids. Here instead we consider a homopolymer model. The ground state is nevertheless found to be highly degenerate. Thus, upon adding sequence heterogeneity, a given sequence has a large predetermined menu of structures to choose its ground state from. Proteins do not have a fixed *θ* angle unlike in our simplified model. The number of amino acids per turn in a protein is approximately 3.6. Here we have a nice integer of 4. There is chiral symmetry breaking in a protein unlike in our model. In fact, the assembly of right-handed and left-handed helices in the PoSSuM phase cannot happen in protein structures. There is also an important difference in the assembly of strands into sheets. The need for close packing promotes the assembly of out of phase strands in the PoSSuM phase, whereas strands making up a protein tend to be in phase. Protein structures are three dimensional and proteins misfold and aggregate into amyloid. The PoSSuM phase, in contrast, is beautifully packed in two layers.

Francis Crick noted [29]: *Physicists are all too apt to look for the wrong sorts of generalizations, to concoct theoretical models that are too neat, too powerful, and too clean. Not surprisingly, these seldom fit well with data*. While our model is, in fact, a too neat and clean model, it does not and nor is it meant to describe proteins, the amazing molecular machines of life. It nevertheless carries important lessons for physics. What we have demonstrated here is the existence of a novel phase of matter in the context of a simplified chain model. The continuous *θ*-transition in standard chain models is replaced by a first order transition. The low temperature PoSSuM phase is characterized by a spontaneous dimensional reduction with the ground states occupying two layers. There are numerous non-trivially related ground states arising from the modular building blocks of helices and strands. One cannot help but wonder what useful hints the PoSSuM phase might offer for understanding the magnificent protein native state structures.

## Acknowledgements

We are indebted to George Rose for collaboration and inspiration. We are very grateful to Pete von Hippel for his warm hospitality and to him and Brian Matthews for stimulating conversations.

## Author Contributions

TŠ and JRB conceived the ideas for the calculations. TŠ carried out the calculations. JRB wrote the paper. All authors participated in understanding the results and reviewing the manuscript.

## References

[1] M. Rubinstein and R. H. Colby, Polymer Physics (Chemistry). Oxford University Press, 1 ed., 2003.

[2] M. Doi and S. F. Edwards, The theory of polymer dynamics. Clarendon Press, New York, 1993.

[3] C. Clementi, A. Maritan and J. Banavar, “Folding, design, and determination of interaction potentials using off-lattice dynamics of model heteropolymers”, Phys. Rev. Lett. 81, 3287–3290 (1998).

[4] J. E. Magee, V. R. Vasquez and L. Lue, “Helical structures from an isotropic homopolymer model”, Phys. Rev. Lett. 96, 207802 (2006).

[5] I. Coluzza, “A Coarse-Grained Approach to Protein Design: Learning from Design to Understand Folding”, PLoS One 6, e20853 (2011).

[6] D. K. Klimov and D. Thirumalai, “Cooperativity in protein folding: from lattice models with side-chains to real proteins”, Folding & Design 3, 127–139, (1998).

[7] M. Muthukumar and C. Y. Kong, “Simulation of polymer translocation through protein channels”, Proc. Natl. Acad. Sci. USA 103, 5273–5278 (2005).

[8] J. R. Banavar, M. Cieplak, T. X. Hoang and A. Maritan, “First-principles design of nanomachines”, Proc. Natl. Acad. Sci. USA 106, 6900–6903 (2009).

[9] T. Škrbić, A. Badasyan, T. X. Hoang, R. Podgornik and A. Giacometti, “From polymers to proteins: the effect of side chains and broken symmetry on the formation of secondary structures within a Wang-Landau approach”, Soft Matter 12, 4783–4793 (2016).

[10] T. Škrbić, T. X. Hoang and A. Giacometti, “Effective stiffness and formation of secondary structures in a protein-like model”. J. Chem. Phys. 145, 084904 (2016).

[11] T. Škrbić, T. X. Hoang, A. Maritan, J. R. Banavar and A. Giacometti, “The elixir phase of chain molecules”, Proteins: Structure, Function, and Bioinformatics 87, 176–184 (2019).

[12] T. Škrbić, T. X. Hoang, A. Maritan, J. R. Banavar and A. Giacometti, “Local symmetry determines the phases of linear chains: a simple model for the self-assembly of peptides”, Soft Matter 15, 5596–5613 (2019).

[13] D. T. Seaton, S. Schnabel, D. P. Landau and M. Bachmann, “From flexible to stiff: Systematic analysis of structural phases for single semiflexible polymers”, Phys. Rev. Lett. 110, 028103 (2013).

[14] T. Škrbić, J. R. Banavar and A. Giacometti, “Chain stiffness bridges conventional polymer and bio-molecular phases”, J. Chem. Phys. 151, 174901 (2019). and references therein

[15] D. Marenduzzo, C. Micheletti, H. Seyed-allaei, A. Trovato and A. Maritan, “Continuum model for polymers with finite thickness”, J. Phys. A: Math. Gen. 38, L277–L283 (2005).

[16] N. C. Fitzkee and G. D. Rose, “Steric restrictions in protein folding: an alpha-helix cannot be followed by a contiguous beta-strand”, Prot. Sci. 13, 633–639 (2004).

[17] G. D. Rose, “In Memoriam: Professor G.N. Ramachandran (1922 - 2001)”, Perspective - Proteins: Structure, Function, and Bioinformatics 10, 1691–1693 (2001).

[18] F. Wang and D. P. Landau, “Efficient, Multiple-Range Random Walk Algorithm to Calculate the Density of States”, Phys. Rev. Lett. 86, 2050 (2001).

[19] M. P. Taylor, W. Paul and K. Binder, “Phase transitions of a single polymer chain: AWang-Landau simulation study”, J. Chem. Phys. 131, 114907 (2009).

[20] M. P. Taylor, W. Paul and K. Binder, “All-or-none proteinlike folding transition of a flexible homopolymer chain”, Phys. Rev. E 79, 050801 (2009).

[21] M. P. Taylor, W. Paul and K. Binder, “On the polymer physics origins of protein folding thermodynamics”, J. Chem. Phys. 145, 174903 (2016).

[22] N. Madras and. D. Sokal, “The pivot algorithm: highly efficient Monte Carlo method for the self-avoiding walk”, J. Stat. Phys. 50, 109–186 (1988).

[23] S. Doniach, T. Garel and H. Orland, “Phase diagram of a semiflexible polymer chain in a theta solvent: Application to protein folding”, J. Chem. Phys. 105, 1601–1608 (1996).

[24] C. Maffi, M. Baiesi, L. Casetti, F. Piazza and P. De Los Rios, “First-order coil-globule transition driven by vibrational entropy”, Nat. Commun. 3, 1065 (2012).

[25] T. E. Creighton, Proteins: structure and molecular properties. New York, Ed. W. H. Freeman, 1993.

[26] I. Bahar, R. L. Jernigan and K. A. Dill, Protein Actions. Garland Science, Taylor & Francis Group, 2017.

[27] A. M. Lesk, Introduction to Protein Science: Architecture, function and genomics. Oxford University Press, 2 ed., 2004.

[28] All-atoms simulations including side-chains have provided invaluable insights into proteins. Please see the three books above for details.

[29] F. Crick, What mad pursuit. Basic Books, New York, 1988.

